# Sex-dependent hypothalamic microinflammation and microglial polarization: The role of JNK in high-fat diet-induced insulin resistance

**DOI:** 10.1101/2025.11.11.687241

**Authors:** Maria Rodriguez-Garcia, Montserrat Bolaños-Hurtado, Aina Redondo, Carme Caelles, Melania González, Elisenda Sanz, Albert Quintana, Sebastian Zagmutt, Rosalia Rodríguez-Rodriguez

**Affiliations:** Department of Biomedical Sciences, Faculty of Medicine and Health Sciences, Universitat Internacional de Catalunya, Sant Cugat del Vallès, 08195, Spain; Institute of Biomedicine of the University of Barcelona (IBUB), Barcelona, Spain; Department of Biochemistry and Physiology, School of Pharmacy and Food Sciences, University of Barcelona, Barcelona, Spain; Institut de Neurociències, Universitat Autònoma de Barcelona, Bellaterra, Spain; Departament de Biologia Cellular, Fisiologia i Immunologia, Universitat Autònoma de Barcelona, Barcelona, Spain; Centro de Investigación Biomédica en Red de Fisiopatología de la Obesidad y la Nutrición (CIBEROBN), Instituto de Salud Carlos III, Madrid, 28029, Spain

**Keywords:** Hypothalamic microinflammation, obesity, insulin resistance, c-Jun N-terminal kinase (JNK), sexual dimorphism

## Abstract

Obesity and its associated metabolic disorders, including insulin resistance and type 2 diabetes, are major global health challenges. Early hypothalamic microinflammation is emerging as a key contributor to the onset of metabolic dysfunction, but its temporal dynamics, underlying mechanisms, and sex-specific differences remain unclear. Here, we examined the effects of short-term high-fat diet (HFD) exposure on hypothalamic microinflammation, glial activation, and insulin signaling in male and female mice.

Males rapidly developed hypothalamic microinflammation, characterized by increased pro-inflammatory cytokines, reactive gliosis, and impaired insulin signaling, whereas females did not show these alterations. Interestingly, we also observed a sex-specific pattern in microglial M2 polarization: females maintained a sustained M2 response throughout the experimental period, while males exhibited only a transient peak that declined in the following days. These sex-specific differences in microglial dynamics and polarization may be linked to the c-Jun N-terminal kinase pathway, a classical mediator of inflammatory signaling.

To further explore this, we analyzed mice lacking JNK3, the CNS-enriched isoform of JNK. Loss of JNK3 disrupted early microglial polarization, abolished the transient M2 response, and predisposed males to enhanced gliosis and partial hypothalamic insulin resistance, while females retained partial resilience. These findings indicate that JNK3 contributes to the regulation of microglial dynamics and polarization in response to metabolic stress. Overall, our study highlights a sex-dependent role of microglia and JNK in shaping early hypothalamic microinflammation and central insulin sensitivity, providing potential targets for intervention in obesity-associated metabolic disease.

## Introduction

Obesity and its related metabolic complications, such as insulin resistance and type 2 diabetes, remain among the most pressing global health challenges of the 21st century (Ng et al., 2025; Pillon et al., 2021). While traditionally considered a disorder of peripheral metabolic tissues such as adipose tissue, liver, and muscle, a growing body of evidence has positioned the hypothalamus, a central regulator of energy balance and glucose homeostasis, as a critical site of early pathological changes in obesity (Argente et al., 2025; Rodríguez-Rodríguez et al., 2023; Timper & Brüning, 2017). This vulnerability is thought to arise from the hypothalamus’ central role in sensing and integrating peripheral nutrient and hormonal signals, making it particularly sensitive to metabolic stress (González-García et al., 2016). Consequently, high-fat diet (HFD) feeding rapidly triggers a state of low-grade inflammation in this region, often preceding overt weight gain or systemic metabolic dysfunction (Bhusal et al., 2021; Cai & Khor, 2019; Thaler et al., 2012; Zagmutt et al., 2025). This hypothalamic microinflammation disrupts neuronal function, impairs hormonal signaling, and contributes to the loss of homeostatic control of energy intake and expenditure (Nampoothiri et al., 2022; Valdearcos et al., 2015). However, the temporal dynamics, molecular mechanisms, and potential sex differences underlying this early inflammatory process remain incompletely understood.

Recent studies have shown that glial cells, particularly microglia and astrocytes, play a pivotal role in the initiation and maintenance of hypothalamic inflammation in response to nutritional excess (Kwon et al., 2017; Sugiyama et al., 2020; Valdearcos et al., 2014). These glial populations respond within hours of HFD exposure, releasing pro-inflammatory cytokines such as interleukin-1β (IL-1β), tumor necrosis factor-α (TNF-α), and others (Cansell et al., 2021). This acute inflammatory response, initially restricted to specific hypothalamic nuclei, is thought to disrupt neuronal metabolic sensing and impair the hypothalamic control of whole-body energy homeostasis.

In addition, interactions between different glial cell types and local neuronal populations may amplify or dampen the inflammatory response, indicating a complex and tightly regulated network that determines the extent of hypothalamic dysfunction. Understanding these interactions is essential to identify potential therapeutic targets for early metabolic disturbances.

Inflammation has long been associated with the development of insulin resistance, a central feature of metabolic disease (Rehman & Akash, 2016). While this relationship has been extensively studied in peripheral tissues, the impact of inflammation on central insulin signaling, particularly in the hypothalamus, remains less clear. The hypothalamus is highly sensitive to insulin, which acts to inhibit food intake, stimulate energy expenditure, and regulate glucose metabolism (Kullmann et al., 2025; Pan et al., 2023; Varela & Horvath, 2012). Impairments in central insulin signaling have been linked to hyperphagia and metabolic imbalance, positioning central insulin resistance as a key node in the pathogenesis of obesity (Chen et al., 2017). Yet, it is still poorly understood whether hypothalamic microinflammation directly contributes to central insulin resistance, and whether this relationship is modulated by sex-specific factors.

The c-Jun N-terminal kinase (JNK) signaling pathway has long been recognized as a crucial intracellular mediator of inflammation and insulin resistance in peripheral metabolic tissues such as liver, muscle, and adipose tissue (Feng et al., 2020; Hirosumi et al., 2002). Activation of JNK promotes serine phosphorylation of insulin receptor substrate-1 (IRS-1), thereby impairing insulin signaling and contributing to metabolic dysfunction (Copps & White, 2012). However, despite extensive evidence in peripheral organs, the role of JNK signaling in hypothalamic microinflammation remains comparatively underexplored. Notably, the isoform predominantly expressed in the brain is JNK3, which appears to be selectively activated in neurons and glial cells in response to metabolic and inflammatory stress (Nogueiras & Sabio, 2021; Priego et al., 2023). Some studies using LPS-induced hypothalamic inflammation suggest that JNK may play an important role in initiating both hypothalamic inflammation and central insulin resistance, but the precise molecular mechanisms involved remain poorly understood (Rorato et al., 2017). Taken together, these observations highlight the hypothalamus as a central node linking nutritional excess, glial activation, and early metabolic dysregulation. However, the molecular mechanisms underlying early hypothalamic microinflammation, including the specific role of JNK, remain incompletely understood. Moreover, potential sex differences in glial responses and microglial polarization have been largely overlooked. This study aims to fill these gaps by investigating the molecular pathways driving early hypothalamic microinflammation and dissecting sex-specific patterns in microglial activation and polarization in response to short-term high-fat diet exposure. By elucidating these mechanisms, we aim to provide a better understanding of the early events that predispose to central insulin resistance and metabolic dysfunction, potentially revealing novel targets for preventive intervention.

## Materials and Methods

### Animals

Male and female C57BL/6J wild-type (WT) mice (Envigo, Spain) and global JNK3 knockout mice (JNK3-KO; kindly provided by Dr. Carme Caelles, Universitat de Barcelona) (Yang et al., 1997) aged 8–10 weeks were used. Both sexes mice were included unless otherwise specified. Animals were maintained under a 12 h light/dark cycle (lights on at 8:00 a.m.) in temperature-and humidity-controlled conditions with ad libitum access to water and chow unless otherwise specified. Mice were group-housed to avoid isolation stress. At the study endpoint, animals were euthanized by cervical dislocation or intracardiac perfusion for tissue collection. All animal procedures were performed in agreement with European guidelines (2010/63/EU) and approved by the Universitat Autònoma de Barcelona and Universitat of Barcelona Local Ethical Committees.

### Dietary interventions

For high-fat diet (HFD) studies, mice were acclimatized for 1 week on normal chow diet (NCD; 10% kcal from fat, D12450J, Research Diets) before switching to HFD (60% kcal from fat, D12492, Research Diets). At the experimental endpoint, mice were fasted for 1 h prior to sacrifice and tissue collection.

### Body weight and food intake

Body weight was measured at end time points and normalized to baseline (day 0). Food intake was assessed by weighing food every 24 h. In selected experiments, automated feeding systems (Smart Waiter Feeder Weigh 90189084, TSE systems, Berlin, Germany) were used to continuously monitor food intake in individually housed mice during NCD or HFD exposure.

### Intracardiac perfusion and brain fixation

Mice were anesthetized with ketamine/xylazine and transcardially perfused with PBS followed by 4% neutral buffered formalin (Sigma-Aldrich). Brains were post-fixed overnight and cryoprotected in 30% sucrose before freezing and storage at −80 °C.

### Insulin tolerance tests

For peripheral insulin tolerance tests (ITT), mice were fasted for 6 h and received intraperitoneal injections of insulin (1.5 U/kg body weight). For central ITT, 100 μU of insulin were administered icv. Blood glucose was measured at baseline and at 15, 30, 60, 90, and 120 minutes post-injection, as described (Fosch et al., 2025).

### Intracerebroventricular administration of insulin

Cannulae were stereotaxically implanted into the lateral ventricle (coordinates relative to Bregma: −0.58 mm AP, +1 mm ML, −2.2 mm DV) under ketamine/xylazine anesthesia. Correct placement was verified by the angiotensin II-induced drinking response. Only responsive animals were included (Rodríguez-Rodríguez et al., 2019). Seven days after surgery, mice were exposed to NCD or HFD for seven more days. On the experimental day, mice were fasted for 6 h and received icv injections of either saline or insulin (100 μU in 2 μL). Brains were collected 1 h later following perfusion.

### pAkt/Akt signaling assay

Mice fasted overnight were anesthetized and injected with insulin (5 U) into the inferior vena cava. Ten minutes later, hypothalamus was collected for Western blot analysis.

### Cell culture

Murine microglial BV2 cells (kindly provided by Dr. Elisenda Sanz, Universitat Autònoma de Barcelona) and hypothalamic neuronal GT1-7 cells (SCC116, Merck) were cultured in high-glucose DMEM supplemented with 10% fetal bovine serum, 1% penicillin/streptomycin, and 1% glutamine. GT1-7 medium additionally contained 1 mM sodium pyruvate. Cells were used between passages 8–30, maintained at 37 °C in 5% CO₂, and routinely tested for mycoplasma.

### BV2–GT1-7 co-culture and palmitate treatment

BV2 and GT1-7 cells were co-cultured at a 1:3 ratio. To mimic HFD conditions, cells were treated with palmitate (150 μM; Sigma-Aldrich) conjugated to fatty acid–free bovine serum albumin (BSA; Merck) for 1, 4 or 6 h. Control cells received unconjugated BSA.

### RT-qPCR and Western blot

Gene expression was assessed by quantitative RT–PCR and protein levels by Western blot. as previously described (Rodríguez-Rodríguez et al., 2019), and brain sections were analyzed by immunofluorescence microscopy (Fosch et al., 2025). Detailed reagent information, including primer sequences and antibody information, are provided in Supplementary materials.

### Immunofluorescence image analysis

All images were acquired as z-stacks and converted into composite projections by averaging pixel intensity across layers. Image analysis was performed using ImageJ software (NIH, USA).

For intensity-based measurements (e.g., GFAP, α-MSH, insulin receptor-β), mean fluorescence intensity was quantified from the selected regions of interest (ROIs). Astrocyte size was determined by measuring the longest axis, while branch complexity was analyzed using the ImageJ skeletonization plugin. For microglial analysis, Iba1⁺ cells were manually counted in Hoechst/Iba1-merged images. Cell size was determined by measuring the average pixel intensity within a standardized ROI. Polarization was assessed by masking Iba1⁺ cells and quantifying CD206 intensity within the mask. For nuclear markers (c-Jun, c-Fos), masks were generated from the Hoechst channel, and fluorescence intensity was quantified within nuclear ROIs. In co-culture experiments, CellBrite labeling was used to define GT1-7 and BV2 cells, and c-Jun intensity was measured within cell-specific masks. Fluorescence intensities were normalized to ROI area to account for differences in cell density. For tissue analysis, at least three slices per mouse were analyzed, with 3–4 mice per group. Two images (left and right hemisphere) were acquired from each region of interest (ARC or PVN). For immunocytochemistry, a minimum of 10 images per slide were collected, with three experimental replicates, and at least three slides per condition in each replicate.

### Statistical analysis

Data are expressed as mean ± SEM. The number of animals or independent replicates per experiment is reported in figure legends. Comparisons between two groups were performed using unpaired two-tailed Student’s t test; multiple groups were analyzed by one-or two-way ANOVA with Tukey’s or Šídák’s post hoc tests, as appropriate. Analyses were performed using Prism v9.0 (GraphPad). P < 0.05 was considered statistically significant.

## Results

### Temporal and Sex-Dependent Modulation of Hypothalamic Inflammation by High-Fat Diet

To deeply explore the dynamics of hypothalamic microinflammation induced by HFD, male and female mice were exposed to a 60% calorie HFD for 1, 3, 7, and 14 days. In males, consistent with previous reports (Morari et al., 2014; Thaler et al., 2012), we observed a progressive increase in the expression of pro-inflammatory cytokines such as IL-6 and TNF-α by day 7 in the hypothalamus, which persisted at day 14 (Fig. 1A). In contrast, females did not exhibit significant upregulation of these pro-inflammatory markers at any time point (Fig. 1C). Interestingly, IL-10, a potent anti-inflammatory cytokine, displayed sex-specific temporal dynamics. In males, IL-10 expression increased significantly by day 7, suggesting a delayed anti-inflammatory response (Fig. 1B). In females, however, IL-10 levels remained unchanged throughout the time course (Fig. 1D). These findings indicate a dynamic yet sex-dependent balance between pro-and anti-inflammatory signaling in the hypothalamus during early HFD exposure.

**Figure 1.**
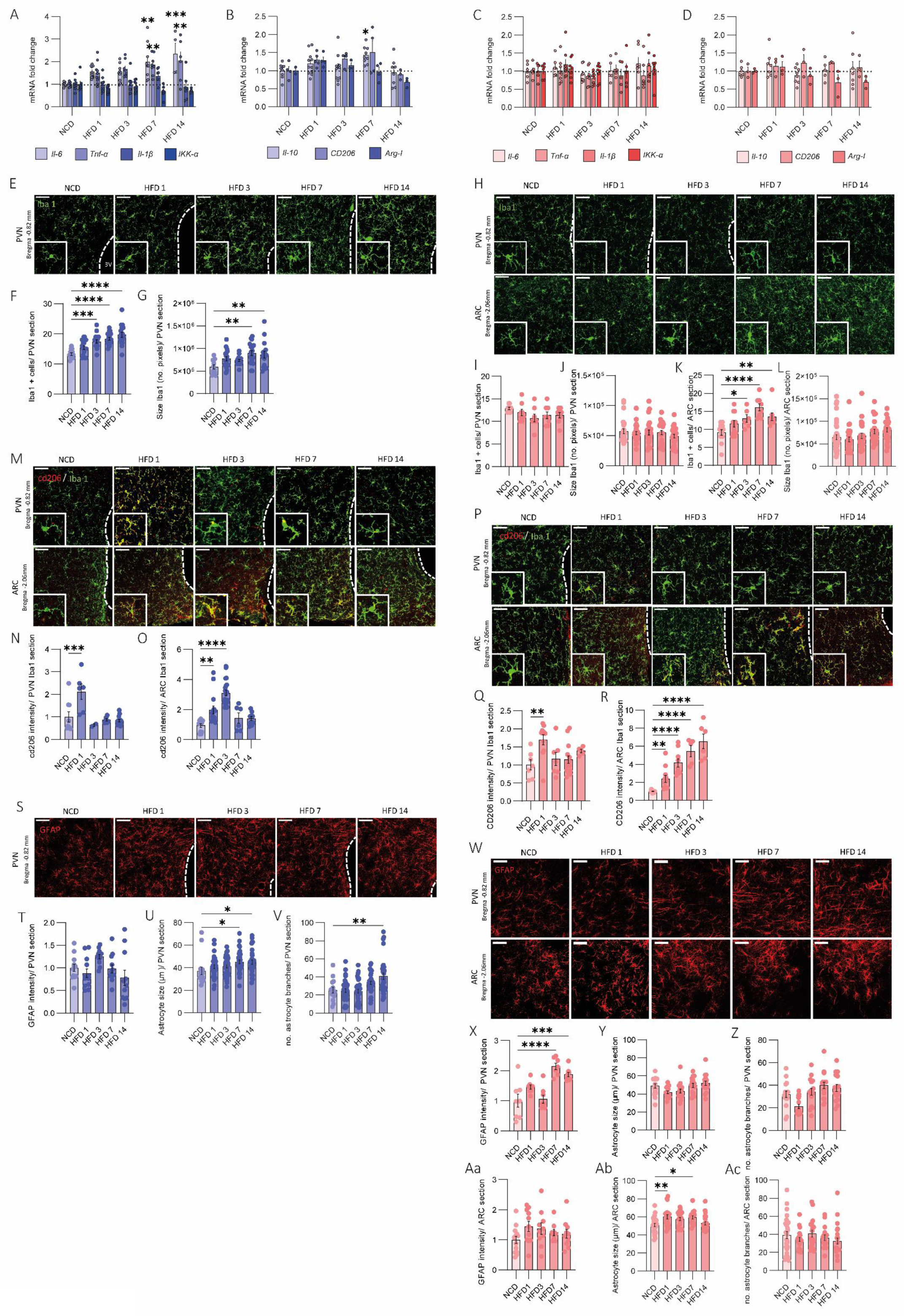
Phenotypic hypothalamic characterization of male and female mice in response to short-term HFD feeding. **(A-D) Hypothalamic cytokine expression**. Analysis of the pro-inflammatory cytokines *Il-6, Tnf-α, Il-1β* and *IKK-α* mRNA levels by qRT-PCR of (A) male and (C) female mice. Analysis of the anti-inflammatory cytokines *Il-10, CD206* and *Arg-1* mRNA levels by qRT-PCR of (B) male and (D) female mice. n=3-9/ group. (E-L) Microglial dynamics. (E, H) Representative confocal images of Iba1 marker (green) in PVN sections of male mice and PVN and ARC sections of female mice. Quantifications of Iba1+ cell number of (F) male and (I, K) female mice. Microglia cell size by analyzing no. of pixels of (G) male and (J, L) female mice. n= 3-4/ group. (M-R) M2 microglia polarization. (M, P) Representative confocal images of CD206 marker (red) in Iba1+ cells (green) in PVN and ARC sections of male and female mice, respectively. CD206 intensity normalized by area of Iba1+ cells of PVN and ARC of (N, O) male and (Q, R) female mice. n= 3-4/ group. (S-Ac) Astrocytic dynamics. (S,W) Representative confocal images of GFAP marker (red) in PVN sections of male mice and PVN and ARC sections of female mice. Quantification of GFAP intensity normalized by the control group of (G) PVN male mice and (X, Aa) PVN and ARC female mice. Quantification of astrocytes cell size in µm of (U) PVN male mice and (Y, Ab) PVN and ARC female mice. Quantification of no. of astrocytes branches of (V) PVN male mice and (Z, Ac) PVN and ARC female mice. n= 3-4/ group. Data were represented as mean ± SEM. Statistical significance was determined by ordinary one-way ANOVA followed by Tukey’s multiple comparisons post-hoc test. *p<0.05, **p<0.01, ***p<0.001, ****p<0.0001 versus NCD group. Scale bar 50 µm. 3V: third ventricle.

Reactive gliosis was assessed by analyzing microglial and astrocytic responses. Iba-1 staining revealed a significant increase in the number and activation of microglia in the paraventricular nucleus (PVN) of male mice from day 3 to day 14 (Fig. 1E-G), consistent with prior observations in the arcuate–median eminence (ARC-ME) for rodents (Thaler et al., 2012). In contrast, female mice did not show significant changes in microglial number or activation in the PVN (Fig. 1H upper panel, I and J). At baseline, no significant differences were observed in the number of microglia in the PVN between male and female mice, indicating comparable initial conditions (Supplementary Fig. 1A). In the ARC-ME, females exhibited an increase in microglial number at days 7 and 14, without evident morphological activation (Fig. 1H lower panel, K and L). Importantly, the baseline microglial counts in the ARC-ME of females closely correlate with those reported by Thaler et al. (2010) in male rats, reinforcing the cross-species and sex-translational relevance of this hypothalamic region in early HFD responses.

To further dissect the nature of the microglial response, we decided to characterize CD206 expression as a marker of M2-like anti-inflammatory microglia. In male mice, our results revealed that one day of HFD triggers a robust M2 response in the PVN, which diminishes with prolonged exposure (Fig. 1M upper panel and N). A similar transient M2 peak was observed in the ARC-ME, lasting 1–3 days before declining (Fig. 1M lower panel and O). In females, CD206 expression followed a different pattern: while levels in the PVN remained comparable to those observed in males (Fig. 1P upper panel and Q) the ARC-ME showed a progressive increase in CD206 expression over time (Fig. 1P lower panel and R), suggesting that sustained M2 polarization in this region may contribute to the female-specific resistance to hypothalamic microinflammation during early HFD exposure.

Astrocytic activation, evaluated through GFAP immunostaining, showed increased astrocyte size and branching in the PVN of male mice at days 7 and 14, consistent with early gliosis (Fig. 1S-V). No such changes were detected in females, suggesting absence of astrocytic activation under similar conditions (Fig. W-Ac). When comparing the basal size of astrocytes in the paraventricular nucleus (PVN) of the hypothalamus, we observed that astrocytes were larger in females than in males (Supplementary Fig. 1B).

Importantly, this early inflammatory response detected mainly in the male hypothalamus preceded peripheral metabolic changes. Body weight and food intake were monitored throughout HFD exposure. While both sexes progressively gained weight by day 14 (Supplementary Fig. 1C and F), HFD-fed animals showed reduced food intake when measured in grams; however, due to the higher caloric density of the HFD, their total caloric intake (kcal) remained significantly elevated compared to controls (Supplementary Fig. 1D, E, G and H). Consistently, hepatic analysis revealed only mild IL-1β expression increases in males, with no significant induction of TNF-α or IL-6 (Supplementary Fig. 1I). Interestingly, we observed a significant increase in IL-10 at days 7 and 14 of HFD (Supplementary Fig. 1J), while hematoxylin–eosin staining of the liver revealed no signs of hepatosteatosis (Supplementary Fig. 1K). In subcutaneous WAT, pro-inflammatory cytokines remained unchanged (Supplementary Fig. 1L); however, macrophage analysis suggested a shift toward the M2 phenotype over the time course, as evidenced by increased Mrc1 expression (Supplementary Fig. 1M).

Overall, these findings reveal a sexually dimorphic early response to HFD: in males, central hypothalamic inflammation precedes peripheral changes, with transient M2 microglial activation followed by delayed IL-10 induction and sustained gliosis, while the liver and subcutaneous WAT mount an early anti-inflammatory response, potentially preparing the tissues to cope with impending fatty acid overload. In contrast, females display minimal gliosis and a prolonged M2 microglial phenotype in the hypothalamus, highlighting a central mechanism that may contribute to their greater metabolic resilience during early HFD exposure.

### Divergent Temporal Patterns of JNK Activation in the Hypothalamus Reveal Sex-Specific Inflammatory Signatures Under High-Fat Diet

Given the central role of the JNK pathway in inflammatory signaling, we investigated its involvement in the hypothalamic response to HFD exposure. JNK activation occurs via phosphorylation at Thr183/Tyr185 residues, leading to downstream modulation of inflammation-related transcription factors such as phospho-c-Jun.

To assess pathway activation, we quantified phosphorylated JNK (p-JNK) levels in the hypothalamus at different time points during HFD exposure. In male mice, one day of HFD led to a marked increase in JNK activation, which declined progressively with continued exposure suggesting a transient, early activation pattern (Fig. 2A-D). Interestingly, when measuring JNK in females, we observed a dual pattern: a significant increase was detected after one day of HFD, followed by a decrease, and then a second rise at day 14 (Fig. 2E-H).

**Figure 2.**
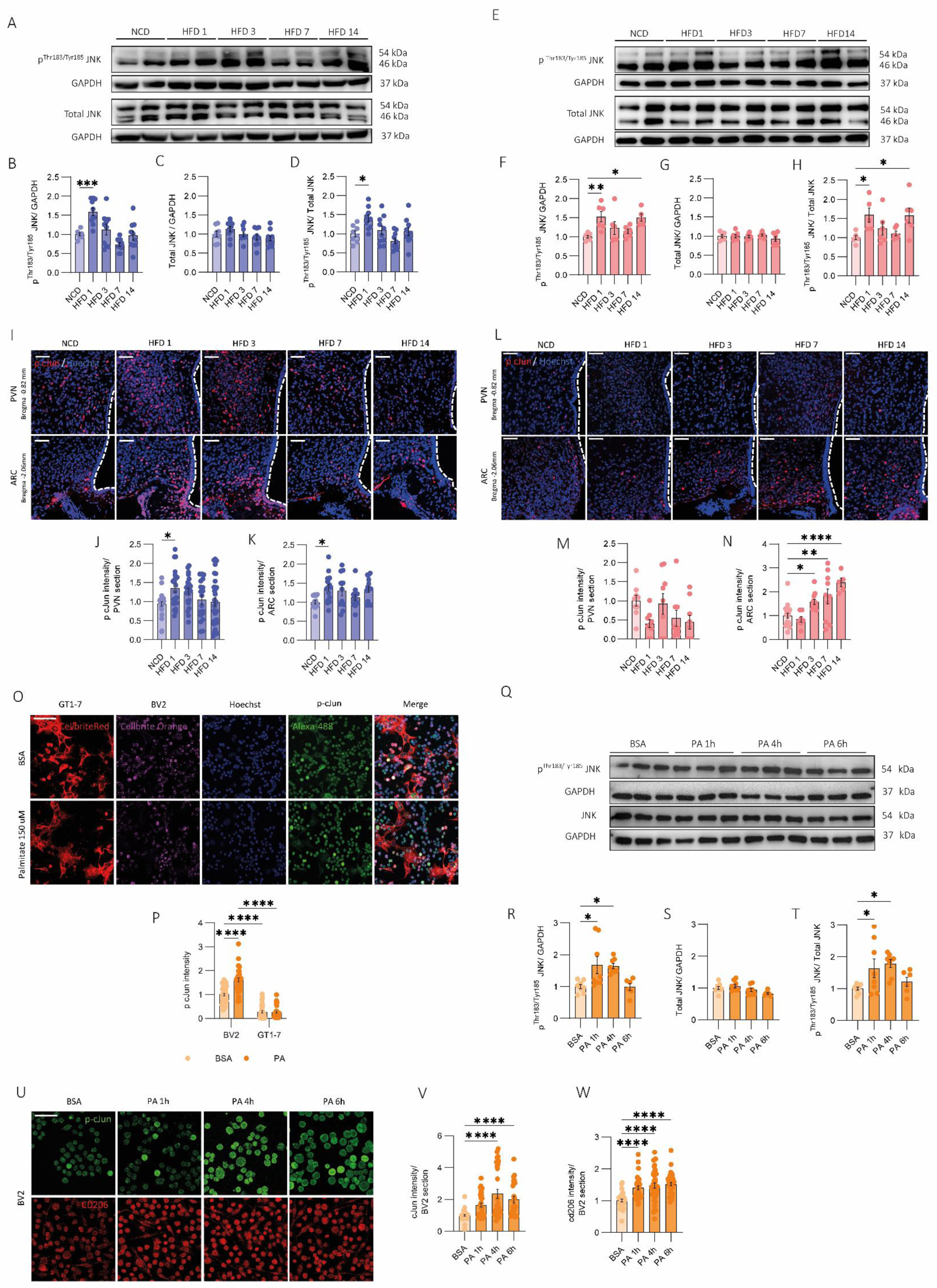
Implication of JNK pathway in hypothalamic microinflammation of male and female mice in response to short-term HFD feeding. (A-H) JNK protein expression in the hypothalamus. Representative Western Blot of JNK protein in the hypothalamus of (A) male and (E) female mice. Quantification of JNK phosphorylation normalized to GAPDH of (B) male and (F) female mice. Quantification of JNK total normalized to GAPDH of (C) male and (G) female mice. Quantification of JNK phosphorylation normalized to total JNK of (D) male and (H) female mice. n= 3-6/ group. (I-N) c-Jun phosphorylation in the PVN and ARC. Representative confocal images of p c-Jun marker (red) and Hoechst marker (blue) in PVN and ARC sections of (I) male and (L) female mice. Quantification of p c-Jun intensity in PVN and ARC of (J, K) male and (M, N) female mice. n= 3-4/ group. (O-P) c-Jun phosphorylation in co-culture of GT1-7 and BV2 after PA exposure. (O) Representative confocal images of p-cJun marker (green), Hoechst (blue), GT1-7 (red cell bright) and BV2 (orange cell bright). (P) Quantification of p-cJun intensity. (Q-T) JNK protein expression in BV2 after PA exposure. (Q) Representative Western Blot of JNK protein in BV2 cells. Quantification of (R) JNK phosphorylation normalized to GAPDH, (S) of JNK total normalized to GAPDH and (T) JNK phosphorylation normalized to JNK total. (U-W) c-Jun phosphorylation and CD206 marker in BV2 after PA exposure. (U) Representative confocal image of p-cJun (green) and CD206 (red) in BV2 cells. Quantification of (V) p-cJun and (W) CD206 intensity in BV2 cells. Data were represented as mean ± SEM. Statistical significance was determined by ordinary one-way ANOVA followed by Tukey’s multiple comparisons post-hoc test. *p<0.05, **p<0.0.01, ***p<0.001, ****p<0.0001. Scale bar 50 µm.

To further validate these findings, we analyzed the spatial distribution of phospho-c-Jun, a direct downstream effector of JNK signaling, in the ARC-ME and PVN through immunodetection. In males, phospho-c-Jun expression followed a similar transient profile, with strong induction at day 1 and a gradual decrease thereafter, consistent with p-JNK dynamics (Fig. 2I-K). Interestingly, in females, phosphorylated c-Jun was detected only in the ARC-ME and increased progressively over time (Fig. 2L–N). These findings indicate that the JNK pathway exhibits a sexually dimorphic and region-specific pattern within the hypothalamus. Interestingly, c-Jun results appear to mirror microglial polarization toward the CD206 phenotype, suggesting a potential functional link between JNK activation and anti-inflammatory microglial responses in a sex-dependent manner, a relationship we further examine in subsequent results.

To further elucidate whether JNK activation plays a critical role in microglial function, we aimed to dissect the cell-type specificity of JNK activation, focusing on microglia as primary sensors of metabolic inflammatory cues. Using an in vitro model, we co-cultured BV2 microglial cells with GT1-7 hypothalamic neurons and exposed them to 150 μM palmitate for 4 hours to mimic HFD-induced stress. Cellbrite Red and Orange dyes were used to identify GT1-7 and BV2 cells, respectively, and phospho-c-Jun immunostaining was performed to assess JNK pathway activation.

Upon direct palmitate exposure, BV2 cells showed a significant and rapid increase in phospho-c-Jun expression, unlike the GT1-7 neuronal cell line, supporting the activation of the JNK pathway specifically in microglia (Fig. 2O and P). In independent cultures, BV2 cells exhibited an early increase in JNK activation after 1 and 4 hours of palmitate incubation (Fig. 2Q-T). Similar results were observed in the analysis of c-Jun in BV2 cells exposed to palmitate, showing a twofold increase in staining after 4 hours of incubation (Fig. 2U upper panel and V), which was more pronounced than in the GT1-7 cells (Supplementary Fig. 2A and B). Importantly, analysis of CD206 in BV2 cells over the same time course paralleled the upregulation of c-Jun under the same conditions (Fig. 2U lower panel and W)

Together, these results suggest that microglia are the early and predominant responders to metabolic stressors such as palmitate, initiating an inflammatory signaling cascade through the JNK pathway. Importantly, both in vivo and in vitro data support a functional link between JNK activation and the regulation of CD206 expression, highlighting a sex-specific mechanism through which microglia shape the differential inflammatory landscape observed during early HFD exposure.

### Early Hypothalamic Microinflammation Drives the Onset of Hypothalamic Insulin Resistance in Male Mice

Inflammation has long been associated with the development of metabolic disorders such as insulin resistance, a link that has been extensively explored in peripheral tissues. However, its impact on central insulin sensitivity, particularly in the hypothalamus, remains less understood. Given the sex-specific differences in hypothalamic inflammatory responses observed in our study, we aimed to investigate whether microinflammation is associated with hypothalamic insulin resistance.

To address this, we focused on the 7-day HFD time point, when male mice exhibited a clear pro-inflammatory phenotype that remained restricted to the hypothalamus, with no evidence of systemic inflammation, while female mice showed no signs of hypothalamic microinflammation. This experimental window allowed us to specifically assess whether central inflammation correlates with impaired hypothalamic insulin signaling in a sex-dependent manner.

We first evaluated the phosphorylation of Akt, a key downstream mediator of insulin signaling, in response to insulin stimulation administered via the inferior cava vein in both male and female mice. Under basal conditions (NCD), insulin effectively activated Akt phosphorylation in the whole hypothalamus of both sexes (Fig. 3A-H). However, after 7 days of HFD, insulin-induced Akt phosphorylation was significantly blunted in males, indicating the development of hypothalamic insulin resistance (Fig. A-D, HFD condition). In contrast, female mice maintained their hypothalamic responsiveness to insulin even after HFD exposure (Fig. 3E-H, HFD condition), suggesting that their resistance to diet-induced inflammation may help preserve central insulin sensitivity.

**Figure 3.**
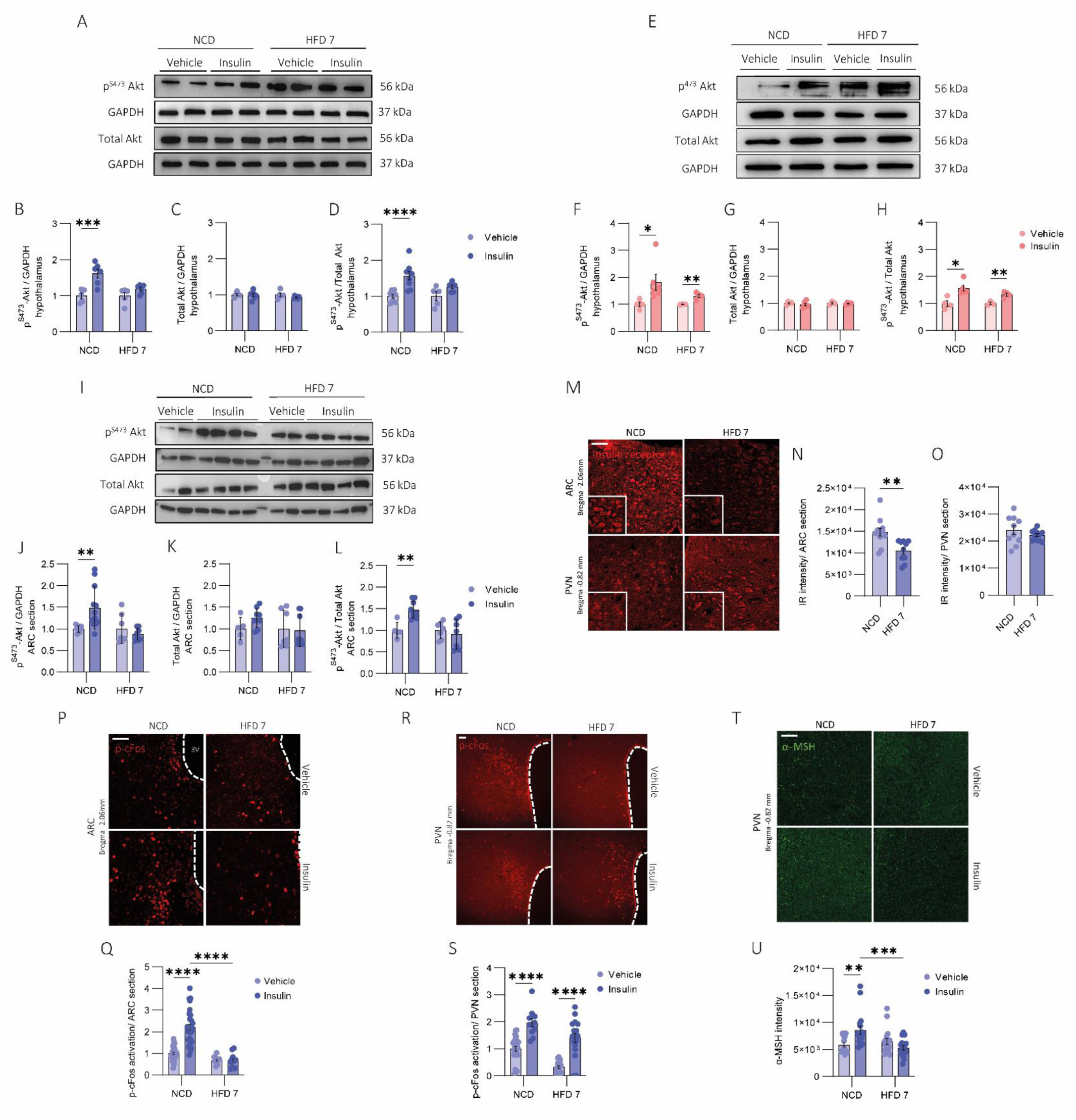
Hypothalamic insulin response of male and female mice after 7 days of HFD feeding. (A-H) Akt hypothalamic activation after insulin administration in male and female mice. Representative Western Blot of Akt protein of the hypothalamus of (A) male and (E) female mice. Quantification of Akt phosphorylation normalized to GAPDH of (B) male and (F) female mice. Quantification of total Akt normalized to GAPDH of (C) male and (G) female mice. Quantification of Akt phosphorylation normalized to total Akt of (D) male and (H) female mice. (I-L) Akt arcuate nucleus activation after insulin administration in male mice. (I) Representative Western Blot of Akt protein in the ARC tissue. Quantification data of (J) Akt phosphorylation normalized to GAPDH, (K) total Akt normalized to GAPDH, and (L) Akt phosphorylation normalized to total Akt. (M-O) Insulin receptor expression in ARC and PVN of male mice. (M) Representative confocal images of insulin receptor marker (red) in ARC and PVN sections. Intensity quantification of insulin receptor in the (N) ARC and in the (O) PVN. (P-S) p-cFos neuronal response to insulin icv administration in ARC and PVN of male mice. Representative confocal images of c-Fos phosphorylation in the (P) ARC and in the (R) PVN sections. Quantification of c-Fos activation in the (Q) ARC and in the (S) PVN. (T-U) α-MSH response to insulin icv administration in PVN of male mice. (T) Representative confocal image in PVN section of male mice of α-MSH (green). (U) Quantification of α-MSH intensity. n= 4-6/ group. Data were represented as mean ± SEM. Statistical significance was determined by t-student or ordinary two-way ANOVA followed by Tukey’s multiple comparisons post-hoc test. *p<0.05, **p<0.0.01, ***p<0.001, ****p<0.0001. Scale bar 50 µm.

Given the development of hypothalamic insulin resistance observed in male mice, we next sought to explore the molecular mechanisms underlying this effect. Since the ARC is a key site for nutrient sensing and has direct contact with the bloodstream through fenestrated capillaries, we repeated the Akt/p-Akt analysis in ARC sections. We observed a similar pattern to that seen in whole hypothalamus: a robust insulin response under basal conditions, which was blunted after 7 days of HFD (Fig. 3I-L). This supports the idea that early hypothalamic microinflammation leads to central insulin resistance without affecting peripheral insulin sensitivity, as confirmed by unchanged insulin tolerance test (ITT) and insulin sensitivity index (kITT) results (Supplementary Fig. 3A-C).

To further investigate how microinflammation might disrupt central insulin action, we quantified insulin receptor (IR) levels in two key hypothalamic nuclei: the ARC and the PVN. After 7 days of HFD, IR expression was markedly reduced in the ARC but remained unchanged in the PVN (Fig. 3M-O), indicating region-specific alterations in insulin signaling components.

We then asked whether this alteration could affect neuronal responsiveness to insulin. To this end, we measured neuronal activation by assessing c-Fos expression following icv insulin administration. As previously described, insulin promotes neuronal activation in both ARC and PVN under normal dietary conditions (Fig. 3P-S, NCD and insulin condition). However, after 7 days of HFD, insulin failed to induce c-Fos expression in the ARC, whereas in the PVN, a partial response was preserved showing increased c-Fos-positive neurons compared to baseline HFD conditions, but not reaching the levels observed under a normal diet (Fig. 3P-S, HFD and insulin condition).

Given the functional connectivity between first-order neurons in the ARC, such as AgRP and POMC neurons, and second-order neurons in the PVN, we also examined the levels of α-MSH, a key neuropeptide released from the ARC to the PVN in response to insulin. As expected, insulin icv administration increased α-MSH levels in the PVN under control conditions. However, this effect was abolished after 7 days of HFD (Fig. 3T and U).

Together, these findings suggest that HFD-induced hypothalamic microinflammation disrupts insulin signaling beginning in the ARC, subsequently impairing downstream neuropeptide communication with other hypothalamic regions such as the PVN. This progression highlights the ARC as an early and vulnerable target of diet-induced central insulin resistance in males, whereas females initially maintain a normal hypothalamic insulin response despite the same dietary challenge, further supporting a sex-specific resilience mechanism during the early stages.

### Loss of JNK3 Contributes to Hypothalamic Microinflammation and Impaired Anti-Inflammatory Response in Male Mice

To further explore the role of JNK pathway activation in microglial dynamics, we focused on the JNK3 isoform, which is specifically expressed in the central nervous system. Using 8-week-old male and female JNK3⁻/⁻ mice (Supplementary Fig. 4A), we first assessed basal levels of inflammatory cytokines in the hypothalamus under a control diet. Our results showed that JNK3-deficient mice exhibited elevated expression of the pro-inflammatory cytokine *Il-1β* in both sexes, compared to their wild-type littermates (Fig. 4A and C). No changes were observed in the anti-inflammatory cytokine *Il-10* (Fig. 4B and D), suggesting that the loss of JNK3 promotes a low-grade pro-inflammatory state without a compensatory anti-inflammatory response.

**Figure 4.**
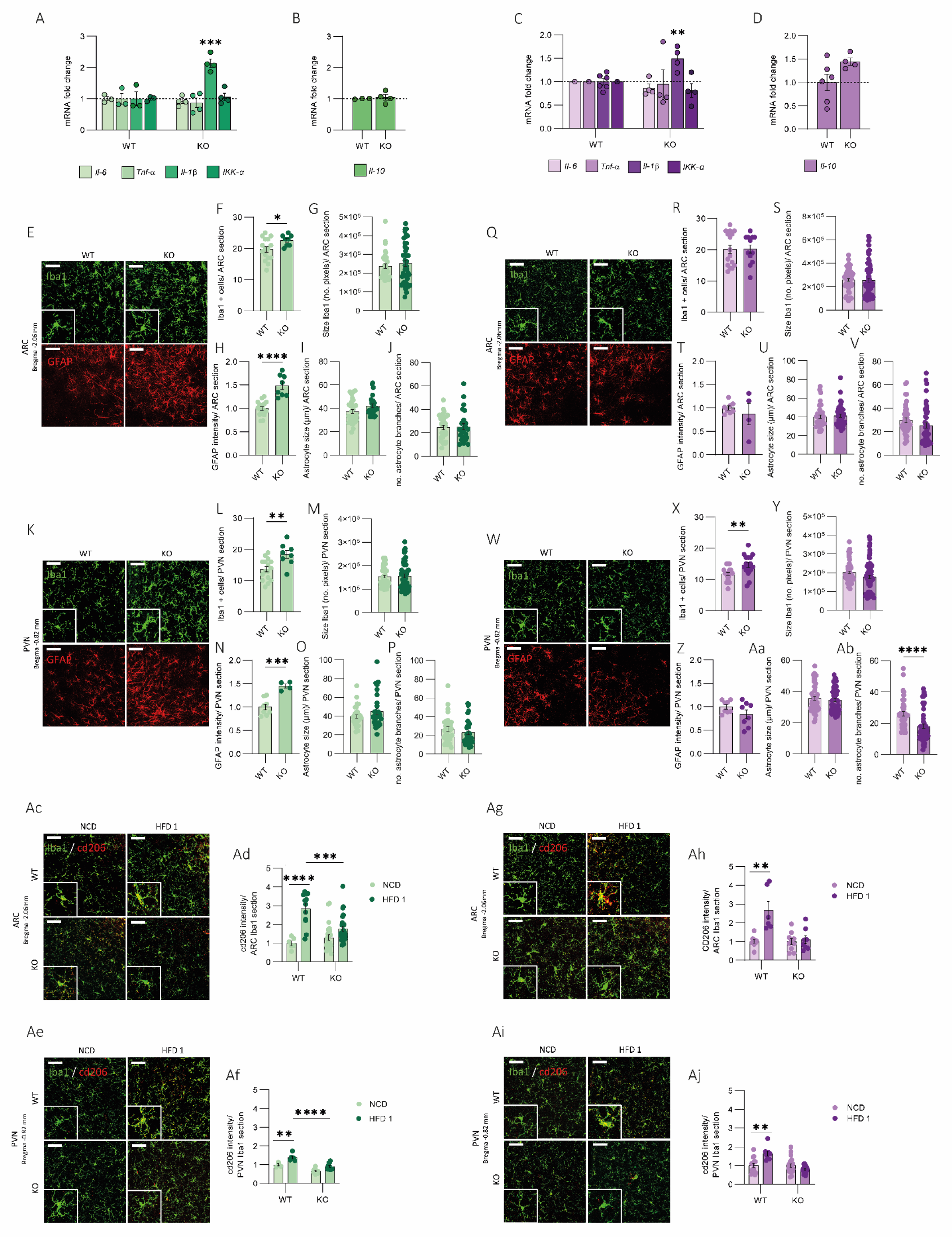
Characterization of JNK3-KO mice. (A-D) Hypothalamic cytokine expression. Analysis of the pro-inflammatory cytokines *Il-6, Tnf-α, Il-1β* and *IKK-α* mRNA levels by qRT-PCR of (A) male and (C) female mice. Analysis of the anti-inflammatory cytokines *Il-10, CD206* and *Arg-1* mRNA levels by qRT-PCR of (B) male and (D) female mice. n=3-6/ group. (E-Ab) Microglial dynamics. (E, K, Q, W) Representative confocal images of Iba1 marker (green) and GFAP marker (red) in ARC and PVN sections of male and female mice. Quantification of Iba1+ cells number of (F) ARC of male, (L) PVN of male, (R) ARC of female and (X) PVN of female. Quantification of microglia cell size by analyzing no. of pixels of (G) ARC of male, (M) PVN of male, (S) ARC of female and (Y) PVN of female. Quantification of GFAP intensity of (H) ARC of male, (N) PVN of male, (T) ARC of female and (Z) PVN of female. Quantification of astrocyte cell size in µm of (I) ARC of male, (O) PVN of male, (U) ARC of female and (Aa) PVN of female. Quantification of no. of branches per cell of (J) ARC of male, (P) PVN of male, (V) ARC of female and (Ab) PVN of female. n= 3/ group. (Ac-Aj) M2 microglia polarization. (Ac, Ag, Ae, Ai) Representative confocal images of CD206 marker (red) in Iba1+ cells (green) in PVN and ARC sections of male and female mice. Quantification of CD206 intensity normalized by area of Iba1+ cells in the (Ad) ARC of male, (Af) PVN of male, (Ah) ARC of female and (Aj) PVN of female. n= 3/ group. Data were represented as mean ± SEM. Statistical significance was determined by t-student and ordinary two-way ANOVA followed by Tukey’s multiple comparisons post-hoc test. *p<0.05, **p<0.0.01, ***p<0.001, ****p<0.0001 versus NCD group. Scale bar 50 µm.

We then analyzed gliosis in the ARC and PVN of the hypothalamus. Specifically, in the ARC of male mice, we observed an increased number of Iba1+ microglia with no changes in soma size (Fig. 4E, upper panel F and G), whereas female mice showed no differences (Fig. 4Q, upper panel R and S). Consistently, GFAP staining in male mice revealed increased intensity per ARC section without changes in astrocytic size or branching (Fig. 4E, lower panel H–J). In contrast, female mice did not display any significant alterations (Fig. 4Q, lower panel T–V). Similar results were observed in PVN sections, where male mice exhibited increased gliosis, reflected by higher Iba1 and GFAP staining (Fig. 4K–P). In contrast, these changes were much less evident in female mice (Fig. 4W–Ab). These results suggest that, despite the absence of JNK3, female mice display a certain resistance to gliosis.

To evaluate whether JNK3 deficiency affects the early M2 polarization response typically observed after HFD exposure (Figure 1), we exposed male and female JNK3⁻/⁻ mice to 1 day of HFD feeding and assessed microglial polarization by measuring CD206 expression in the ARC and PVN.

In wild-type controls, CD206 expression was upregulated after acute HFD exposure, consistent with a transient anti-inflammatory phenotype described previously. However, this response was abolished in JNK3⁻/⁻ mice of both sexes (Fig. 4Ac-Aj), indicating that JNK3 is required for the microglial shift toward an anti-inflammatory state upon early nutritional stress. Similarly to what we did with the WT animals, we decided to perform the 7-day HFD challenge in the KO mice. Interestingly, this was accompanied by hyperphagia in both males and females, without significant changes in body weight (Supplementary Fig. B-G). This dissociation between food intake and weight gain suggests a potential increase in energy expenditure in the absence of JNK3, raising the possibility of a compensatory thermogenic mechanism.

Analysis of gliosis in KO animals exposed to 7 days of HFD revealed that, in the ARC nucleus of males, both the number and size of glial cells further increased compared to basal conditions, with no changes in astrocytes (Supplementary Fig. 4H–M). In females, gliosis remained at basal levels with no changes in either cell number or size, showing only an increase in astrocytic size after 7 days of HFD (Supplementary Fig. 4T–Y). In contrast, no statistically significant differences were observed in the PVN after 7 days of HFD in either males or females (Supplementary Fig. N–Ae).

Taken together, these findings indicate that JNK3 deficiency predisposes the hypothalamus to a pro-inflammatory state and abolishes the early microglial shift toward the anti-inflammatory CD206⁺ phenotype observed in wild-type mice, affecting both males and females. Notably, this is accompanied by a pronounced sexual dimorphism in gliosis: males exhibit an accelerated glial response both at baseline and following HFD exposure, whereas females retain partial resilience, implying intrinsic protective mechanisms that restrain glial overactivation and contribute to sex-specific regulation of hypothalamic inflammation and energy homeostasis.

### Loss of JNK3 Impairs Central Insulin Signaling in a Sex-Specific Manner

Given the critical role of the hypothalamus in regulating systemic metabolism, we asked whether impaired microglial polarization and inflammation induced by JNK3 deficiency affect central insulin sensitivity. We assessed hypothalamic insulin signaling by measuring Akt phosphorylation following insulin administration via the inferior vena cava. In male JNK3⁻/⁻ mice, insulin failed to induce Akt phosphorylation (Fig. 5A–D), indicating hypothalamic insulin resistance under basal dietary conditions. In contrast, female JNK3⁻/⁻ mice retained insulin responsiveness (Fig. 5E–H), suggesting a sex-specific compensatory mechanism that preserves central insulin signaling.

**Figure 5.**
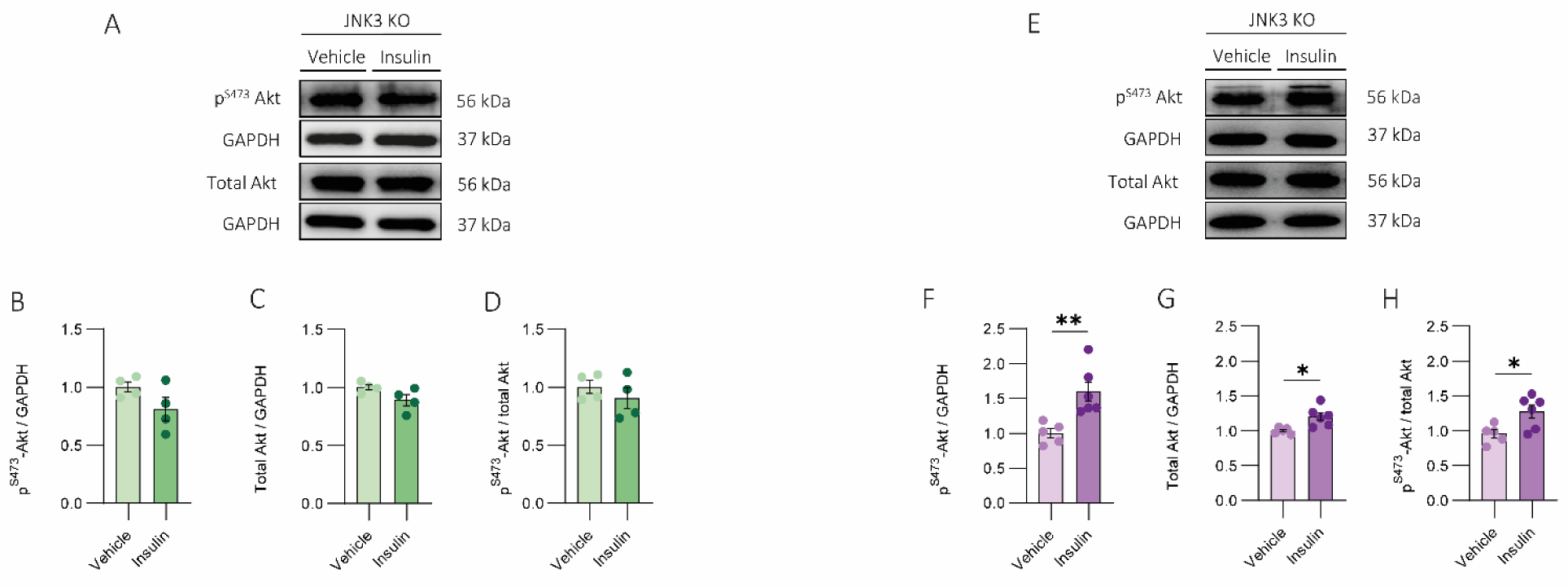
Hypothalamic insulin response of JNK3 KO male and female mice assessed by Akt test. Representative Western Blot of Akt protein of the hypothalamus of (A) male and (E) female mice. Quantification of Akt phosphorylation normalized to GAPDH in (B) male and (F) female mice. Quantification of total Akt normalized to GAPDH in (C) males and (G) female mice. Quantification of Akt phosphorylation normalized to total Akt in (D) males and (H) female mice. Data were represented as mean ± SEM. Statistical significance was determined by t-student. *p<0.05, **p<0.0.01.

Collectively, these results demonstrate that JNK3 is essential for hypothalamic immune homeostasis and central insulin sensitivity. Loss of JNK3 abolishes early CD206⁺ microglial polarization in both sexes, likely predisposing the hypothalamus to a partial pro-inflammatory state. However, males show accelerated gliosis and impaired Akt phosphorylation, whereas females retain partial resilience, maintaining controlled glial activation and preserving hypothalamic insulin signaling. These findings highlight a sex-specific role of JNK3 in linking microglial dynamics to nutrient-sensing pathways.

## Discussion

Our findings identify early hypothalamic microinflammation as a sexually dimorphic process that emerges within days of HFD exposure. Rather than a uniform response, males and females engage distinct glial programs that shape how the hypothalamus senses and adapts to metabolic stress. In line with previous studies showing rapid microglial and astrocytic activation in the ARC of male mice after only three days of HFD exposure (Thaler et al., 2012), our data confirm that males display an accelerated and robust glial response, particularly in the ARC and PVN. In contrast, females exhibit a delayed and attenuated activation, with increased numbers of Iba1⁺ cells in the ARC only after seven days of HFD exposure and no detectable activation in the PVN. These findings indicate that glial reactivity follows a sex-dependent temporal pattern, where males mount a stronger pro-inflammatory response, while females show partial resilience characterized by more protective, tissue-repair-oriented phenotypes (Guneykaya et al., 2018; Han et al., 2021; Lynch, 2022). This resilience may be supported by the greater metabolic flexibility of female glial cells, which allows more efficient utilization of free fatty acids (FFAs) and consequently lower accumulation of these pro-inflammatory substrates in both blood and brain (Cutugno et al., 2024). Altogether, these results position the ARC as an early and vulnerable target of diet-induced inflammation, likely due to its direct exposure to circulating nutrients at the blood–brain barrier, which may transmit metabolic stress signals to other hypothalamic regions, including the PVN (Jais & Brüning, 2022; Steuernagel et al., 2022).

In line with the pro-inflammatory pattern observed in males, previous studies have shown that short-term HFD exposure increases pro-inflammatory cytokine expression in the hypothalamus of male rodents (De Paula et al., 2024; Thaler et al., 2012). Consistently, our results revealed that male mice exhibit elevated hypothalamic pro-inflammatory cytokines from day 7 to day 14, primarily driven by Il-6 and Tnf-α, whereas no activation was detected at 1 or 3 days of HFD. Interestingly, we also observed an acute peak of the anti-inflammatory cytokine Il-10 at day 7. The simultaneous presence of pro-and anti-inflammatory cytokines likely reflects the dynamic nature of microglial activation, whereby microglia do not adopt specific phenotypes but adjust their polarization according to environmental cues, potentially mitigating HFD-induced hypothalamic damage (Guo et al., 2022). This dynamic microglial behavior may reconcile apparent discrepancies between our findings and those of Thaler et al. (2012), who detected a pattern of on-off-on pro-inflammatory activation.

Given these observations, we next explored microglial polarization across sexes and hypothalamic nuclei. The anti-inflammatory capacity is known to differ between sexes, with females generally exhibiting a stronger and more sustained response (Guneykaya et al., 2018). In males, microglia in the ARC and PVN initially polarized toward M2 phenotype after 1 day of HFD exposure, but this polarization rapidly shifted to a M1 phenotype from days 3 to 14, correlating with increased pro-inflammatory cytokine expression. In contrast, female microglia displayed distinct dynamics. In the PVN, activation remained low, with only a transient M2 polarization at 24 hours that quickly subsided, without inducing a pro-inflammatory phenotype, indicating minimal gliosis. In the ARC, M2 polarization occurred at 24 hours and progressively increased over the 14-day exposure period. These data reveal a previously unrecognized sexual dimorphism: female microglia remain predominantly anti-inflammatory or quiescent, whereas male microglia adopt a pro-inflammatory phenotype characterized by gliosis and elevated cytokine expression. These differences may, at least in part, be mediated by hormone-dependent regulation of immune genes, as estrogen has been shown to modulate cytokine production and promote anti-inflammatory polarization at high concentrations (De Paula et al., 2024). Collectively, our findings uncover sex-specific molecular mechanisms governing hypothalamic responses to short-term HFD exposure. In this context, females appear to adapt more efficiently to the initial lipid overload imposed by HFD, mounting protective anti-inflammatory and metabolic responses that may contribute to their premenopausal resilience against obesity and its related comorbidities.

Furthermore, these results address a key question: whether hypothalamic microinflammation in response to HFD precedes the effects of these nutrients on metabolically active peripheral organs such as the liver and white adipose tissue (WAT). Our data confirms that the initial response arises in the hypothalamus, suggesting that long-term alterations in peripheral tissues may be secondary to dysregulation of this central control region. These observations and the temporal dynamics of hypothalamic microinflammation align with the three-phase framework recently proposed by our group (Zagmutt et al., 2025), which conceptualizes the progression of hypothalamic microinflammation into: (1) the Initiation spark, characterized by rapid inflammatory signaling; (2) the Adaptive transition, during which compensatory mechanisms attempt to restore homeostasis; and (3) the Dysfunctional phase, leading to chronic inflammation and metabolic impairment. Our current findings fit within this framework, providing experimental evidence that the early hypothalamic response to HFD represents the initiation stage of this process, preceding peripheral inflammation and insulin resistance.

Consistent with this model, early hypothalamic microinflammation in male mice was detectable before any significant inflammatory changes in the liver or WAT, consistent with previous reports (Thaler et al., 2012). Notably, we also observed early adaptive responses in peripheral tissues, including the infiltration of anti-inflammatory macrophages in WAT and increased Il-10 expression in the liver at days 7 and 14, potentially reflecting a protective mechanism preparing the organism for lipid overload. With prolonged HFD exposure, these compensatory mechanisms may be overwhelmed, ultimately resulting in sustained pro-inflammatory states in peripheral organs (Maric et al., 2014), suggesting that early hypothalamic inflammation may act as a central trigger for downstream metabolic dysregulation.

After years of research, JNK has emerged as a key effector in inflammatory and stress responses and has been widely implicated in the development of insulin resistance (Garg et al., 2021; Nogueiras & Sabio, 2021; Schriever et al., 2020). Our analysis of hypothalamic JNK dynamics revealed that this pathway is also involved in the early response to HFD.

Male mice exhibited a peak in JNK activation 24 hours after HFD exposure, which was transduced via c-Jun phosphorylation in both the ARC and PVN. Notably, this peak coincided with microglial polarization toward the anti-inflammatory M2 phenotype. As JNK and c-Jun were dephosphorylated, M2 polarization declined, permitting the onset of a pro-inflammatory response. A similar temporal relationship was observed in females: c-Jun phosphorylation in the ARC began at day 3 and increased until day 14, paralleling CD206 expression, whereas the PVN remained largely non-reactive, consistent with minimal microglial activation. The alignment between JNK activation and M2 polarization in both sexes suggests a mechanistic link. Supporting this, co-culture experiments with microglia and hypothalamic neurons exposed to palmitic acid revealed that microglia exhibited robust c-Jun phosphorylation and M2 polarization, suggesting that they may represent the primary brain cell type responding to JNK signaling during early metabolic stress.

Although JNK is often described as pro-inflammatory and associated with insulin resistance during long-term HFD exposure, particularly through JNK1/2 isoforms (Nogueiras & Sabio, 2021), our findings suggest that JNK3 may play a distinct role. The three isoforms coexist and appear to require a balanced interaction for proper regulation. Under physiological conditions, JNK3 seems to be activated in states of satiety, where it modulates anorexigenic neurons, in contrast to JNK1/2, would promote increased food intake and decreased metabolism (Garg et al., 2021; Mazzoli et al., 2020; Zeke et al., 2016).

During short-term HFD exposure, JNK3 appears to predominate, inducing M2 polarization and establishing an anti-inflammatory environment that likely acts as a protective mechanism against early hypothalamic stress induced by the diet. Despite this protective role of JNK3, males exhibited only a transient response: M2 polarization occurred after 1 day of HFD but rapidly declined with continued feeding, potentially contributing to hypothalamic microinflammation and increasing susceptibility to metabolic dysregulation. In contrast, females maintained a sustained M2 response throughout the 14-day HFD period, which may underlie their resilience to HFD-induced hypothalamic inflammation and associated metabolic comorbidities. These findings highlight a sex-specific mechanism in which prolonged maintenance of M2 microglia, particularly in the ARC, may confer protection against the metabolic disruptions initiated by HFD.

Prolonged inflammation resulting from chronic HFD feeding is a well-recognized driver of peripheral insulin resistance (Cutugno et al., 2024). However, whether the early hypothalamic microinflammation induced by short-term HFD is sufficient to alter insulin sensitivity remains less understood.

Previous studies have shown that central insulin resistance can arise rapidly: in male rats, just three days of HFD were sufficient to abolish the anorexigenic effects of insulin icv administration (Clegg et al., 2011; Ono, 2019). Consistent with these findings, our results demonstrate that male mice developed hypothalamic insulin resistance after 7 days of HFD. Specifically, we observed reduced IR levels and diminished neuronal responsiveness to insulin stimulation in the ARC, resulting in impaired insulin sensitivity as measured by pAkt/Akt signaling (Ramírez et al., 2022). Given the ARC’s location at the base of the hypothalamus, where the BBB is more permeable (Jais & Brüning, 2022), it is particularly vulnerable to circulating nutrients. The impaired insulin response in the ARC likely disrupted communication with second-order neurons in the PVN, leading to reduced insulin responsiveness in the PVN despite preserved receptor density.

Hypothalamic insulin signaling also regulates hepatic glucose production, and disruption of this pathway after only a few days of HFD exposure has been shown to impair liver function. In contrast, insulin resistance in peripheral tissues such as skeletal muscle and adipose tissue generally emerges after several weeks of HFD feeding, supporting the notion that hypothalamic dysfunction precedes and potentially drives the onset of peripheral insulin resistance.

Importantly, female mice did not develop hypothalamic insulin resistance after 7 days of HFD exposure. Even after 14 days, females maintained JNK activation associated with sustained M2 microglial polarization, thereby preserving an anti-inflammatory hypothalamic state. These findings suggest that the sustained JNK activity and anti-inflammatory milieu in females help safeguard central insulin signaling, potentially contributing to their enhanced capacity to withstand early metabolic stress induced by HFD.

Specifically, JNK3 isoform, which is predominantly expressed in the brain, has been implicated in metabolic protection under HFD conditions, preventing the development of insulin resistance and obesity (Nogueiras & Sabio, 2021). In line with these observations, our data identify JNK3 as a key molecular component underlying the sexual dimorphism in metabolic protection. Beyond its role in energy balance, JNK3 has also been linked to neuronal stress responses, where it exerts pro-apoptotic functions to eliminate damaged neurons (Mazzoli et al., 2020). Loss of this protective mechanism may facilitate chronic neuroinflammation, consistent with our finding that JNK3 deficient mice of both sexes displayed elevated *Il-1β* expression without compensatory changes in the anti-inflammatory cytokine *Il-10*. These results indicate that JNK3 deficiency is sufficient to establish a basal pro-inflammatory state independently of HFD exposure.

This inflammatory environment is closely associated with gliosis (Dorfman & Thaler, 2015), as observed in our models. Male JNK3⁻/⁻ mice exhibited increased numbers of microglia and astrocytes in both the ARC and PVN, without concurrent changes in cell size or morphology, a pattern that may reflect infiltration of peripheral macrophages rather than overactivation of resident glial cells (Mukherjee et al., 2023). Female JNK3⁻/⁻ mice also showed elevated microglial numbers in the PVN, although responses in the ARC were more heterogeneous. Collectively, these findings suggest that JNK3 deficiency promotes basal neuroinflammation and gliosis in both sexes, effectively diminishing the sexual dimorphism observed in wild-type mice.

Upon HFD exposure, gliosis displayed sex-dependent differences in JNK3-/-mice. Males showed an increase in both number and size of microglial cells in the ARC, but not in the PVN, after 7 days of feeding. This regional specificity is consistent with the anatomical position of the ARC, which is directly exposed to circulating factors (Jais & Brüning, 2022). In contrast, astrocytes did not exhibit additional changes in either nucleus, suggesting that HFD-induced gliosis was only modestly superimposed on the already elevated baseline inflammation caused by JNK3 deficiency. Conversely, female JNK3-/-mice failed to develop HFD-induced gliosis in either nucleus, highlighting a preserved resistance to diet-induced neuroinflammation. Thus, while JNK3 loss abolishes sex differences under basal conditions, females retain a unique protection against HFD-induced gliosis, suggesting that additional, JNK3-independent mechanisms contribute to their resilience.

Finally, we demonstrated that JNK3 is required for the polarization of microglia towards the anti-inflammatory M2 phenotype. In its absence, neither male nor female mice exhibited M2 microglial polarization, providing a mechanistic link between JNK3 deficiency, the establishment of a pro-inflammatory status, and the subsequent development of sex-specific insulin resistance.

In line with our previous results regarding the extra protection of the female JNK3-/-mice, our data reveal a striking sex difference in the impact of JNK3 deficiency on hypothalamic insulin signaling. Male JNK3-/-mice displayed insulin resistance, as indicated by reduced pAkt/Akt ratios, consistent with the failure of microglia to polarize towards an anti-inflammatory M2 state and the consequent pro-inflammatory basal state. In contrast, female JNK3-/-mice maintained insulin responsiveness despite the loss of M2 polarization. This divergence suggests that, while both sexes share impaired microglial polarization, females can recruit compensatory mechanisms that preserve hypothalamic insulin sensitivity and confer an additional layer of metabolic protection.

Sex hormones likely contribute to this resilience. Estrogens are key regulators of glucose homeostasis, acting centrally and peripherally, and estrogen deficiency or impaired estrogen receptor signaling is strongly associated with insulin resistance and metabolic dysfunction (Yan et al., 2019). Within the brain, estrogen receptor activation has been shown to influence insulin signaling pathways and promote glucose uptake, with ERα/ERβ directly engaging components of the insulin cascade (Barros & Gustafsson, 2011). Such mechanisms may underlie the preserved insulin response in JNK3-/-females. Nonetheless, hormonal regulation alone is unlikely to fully account for the observed phenotype. Additional, yet unidentified, protective pathways may operate in females, highlighting a critical gap in knowledge given that female hypothalamic insulin signaling remains far less characterized than that of males. In summary, our findings identify JNK3 as a central regulator of hypothalamic inflammation and metabolic responses, with sex-dependent outcomes. While JNK3 deficiency induces basal neuroinflammation in both sexes, its consequences are more pronounced in males, who exhibit both basal insulin resistance and diet-induced microglial activation. In contrast, females show partial protection, being less responsive to HFD-induced gliosis and retaining hypothalamic insulin sensitivity despite the loss of JNK3. These observations underscore the necessity of incorporating sex as a biological variable in metabolic research and point to JNK3 as a potential therapeutic target. Future studies should dissect the molecular mechanisms underlying female resilience, which could reveal novel protective pathways against metabolic dysfunction.

### Conclusions

This study provides new insights into the role of JNK3 in hypothalamic inflammation and insulin resistance, establishing sex-specific differences in the response to HFD. Moving forward, it will be essential to determine whether diet-induced alterations are reversible upon return to a normal chow diet, and to further dissect the molecular pathways that mediate sexual dimorphism in metabolic regulation.

Our results support the notion that the hypothalamus acts as an early driver of peripheral insulin resistance, preceding adipose tissue expansion and systemic metabolic deterioration. This challenges the classical paradigm that insulin resistance arises solely as a secondary consequence of obesity, instead positioning central mechanisms as primary contributors to disease onset. In this light, therapeutic strategies should be revisited to identify novel molecular targets and to incorporate sex-specific considerations.

Importantly, female models must be more systematically investigated to uncover protective mechanisms that may inform precision medicine approaches. Taken together, our findings emphasize the need to move beyond dogmatic frameworks in the field, adopting a more integrative perspective on the origin and progression of metabolic diseases, with the ultimate goal of developing tailored and effective interventions.

## Acknowledgements

This study was supported by the Spanish Ministry of Science and Innovation (MCIN) (PID2020-114953RB-C22 and PID2023-146716OB-I00 to RRR) co-funded by the European Regional Development Fund, the Biomedical Research Centre in Pathophysiology of Obesity and Nutrition (CIBEROBN).

## Notes

### Competing Interest Statement

The authors have declared no competing interest.

