## Supplemental tables and figures for "Sex-dependent hypothalamic microinflammation and microglial polarization: The role of JNK in high-fat diet-induced insulin resistance"

**Supplementary Table 1 |** Primer sequences used for qRT–PCR.

| Gene | Sequence (5' – 3') |
|---|---|
| Il-6 Forward | CTGCAAGAGACTTCCATCCAGT |
| Il-6 Reverse | GAAGTAGGGAAGGCCGTGG |
| Tnf-α Forward | CCAGTGTGGGAAGCTGTCTT |
| Tnf-α Reverse | AAGCAAAAGAGGAGGCAACA |
| Il-1β Forward | CACTACAGGCTCCGAGATGAACAAC |
| Il-1β Reverse | TGTCGTTGCTTGGTTCTCCTTGTAC |
| IKK-α Forward | TGCCTGGCCAGTGTAGCAGTCTT |
| IKK-α Reverse | CAAAGTCACCAAGTGCTCCACGAT |
| Il-10 Forward | GCTCTTACTGACTGGCATGAG |
| Il-10 Reverse | CGCAGCTCTAGGAGCATGTG |
| Plin2 Forward | CCCTCGATTTCAACGTACCC |
| Plin2 Reverse | GCAGCCTGTGGCAATTCA |
| Itgax Forward | GCCATTGAGGGCACAGAGA |
| Itgax Reverse | GAAGCCCTCCTGGGACATCT |
| Mrc1 Forward | CGGTGAACCAAATAATTACCAAAAT |
| Mrc1 Reverse | GTGGAGCAGGTGTGGGCT |
| AgRP Forward | TTTGTCCTCTGAAGCTGTATGC |
| AgRP Reverse | GCATGAGGTGCCTCCCTA |
| Oxytocin Forward | TCTCGCTTGCTGCCTGCTTGG |
| Oxytocin Reverse | GGGAGACACTTGCGCATATCCAG |
| Hprt Forward | GGACCTCTCGAAGTGTTGGATAC |
| Hprt Reverse | GCTCATCTTAGGCTTTGTATTTGGCT |
| CRH Forward | AGGGAGGAGAAGAGAGCGCCCC |
| CRH Reverse | TGCAAGGCAGGCAGGACGAC |

**Supplementary Table 2 |** Antibodies used for Western blot.

| Antibody | Reference | Dilution | Host |
|---|---|---|---|
| Phospho-SAPK/JNK (Thr183/Tyr185) | 9251, Cell Signaling, Danvers, USA | 1:1000 | Rabbit |
| JNK1/2/3 | 9252, Cell Signaling, Danvers, USA | 1:1000 | Rabbit |
| JNK3 | 2305, Cell Signaling, Danvers, USA | 1:1000 | Rabbit |
| Phospho-Akt (Ser473) | 9271 Cell Signaling, Danvers, USA | 1:1000 | Rabbit |
| Akt | 4691 Cell Signaling, Danvers, USA | 1:1000 | Rabbit |
| GAPDH | AM4300, Thermo Fisher, Waltham, USA | 1:50000 | Mouse |
| Anti-Rabbit – HRP | 111-035-144, Jackson, West Grove, USA | 1:10000 | Goat |
| Anti-mouse – HRP | 515-035-003 Jackson, West Grove, USA | 1:20000 | Goat |

**Supplementary Table 3 |** Antibodies used for immunofluorescence.

| Antibody | Reference | Dilution | Host |
|---|---|---|---|
| Iba1 | 019-19741, Fujifilm Wako, Richmond, USA | 1:500 | Rabbit |
| GFAP | 13-0300, Thermo Fisher, Waltham, USA | 1:1000 | Rat |
| CD-206 | 58986, Santa Cruz, Heidelberg, Germany | 1:50 | Mouse |
| c-Jun | 9261, Cell Signaling, Danvers, USA | 1:100 | Rabbit |
| Insulin receptor β | 57342, Santa Cruz, Heidelberg, Germany | 1:50 | Mouse |
| c-Fos | 2250, Cell Signaling, Danvers, USA | 1:200 | Rabbit |
| c-Fos | TA346914, Quimigen, Madrid, Spain | 1:50 | Mouse |
| CRH | NBP3-15356, Novus Biologicals, Madrid, Spain | 1:100 | Rabbit |
| α-MSH | AB5087, Sigma-Aldrich, Madrid, Spain | 1:2000 | Sheep |
| Anti-rabbit Alexa Fluor 488 | a11008 Invitrogen, Barcelona, Spain | 1:1000 | Goat |
| Anti-rabbit Alexa Fluor 568 | a11011, Invitrogen, Barcelona, Spain | 1:1000 | Goat |
| Anti-mouse Alexa Fluor 568 | a11004, Invitrogen, Barcelona, Spain | 1:1000 | Goat |
| Anti-Sheep Alexa Fluor 488 | A-11015, Thermo Fisher, Waltham, USA | 1:1000 | Donkey |
| Anti-rat Alexa Fluor 568 | ab175476, Abcam, Amsterdam, Netherlands | 1:1000 | Goat |

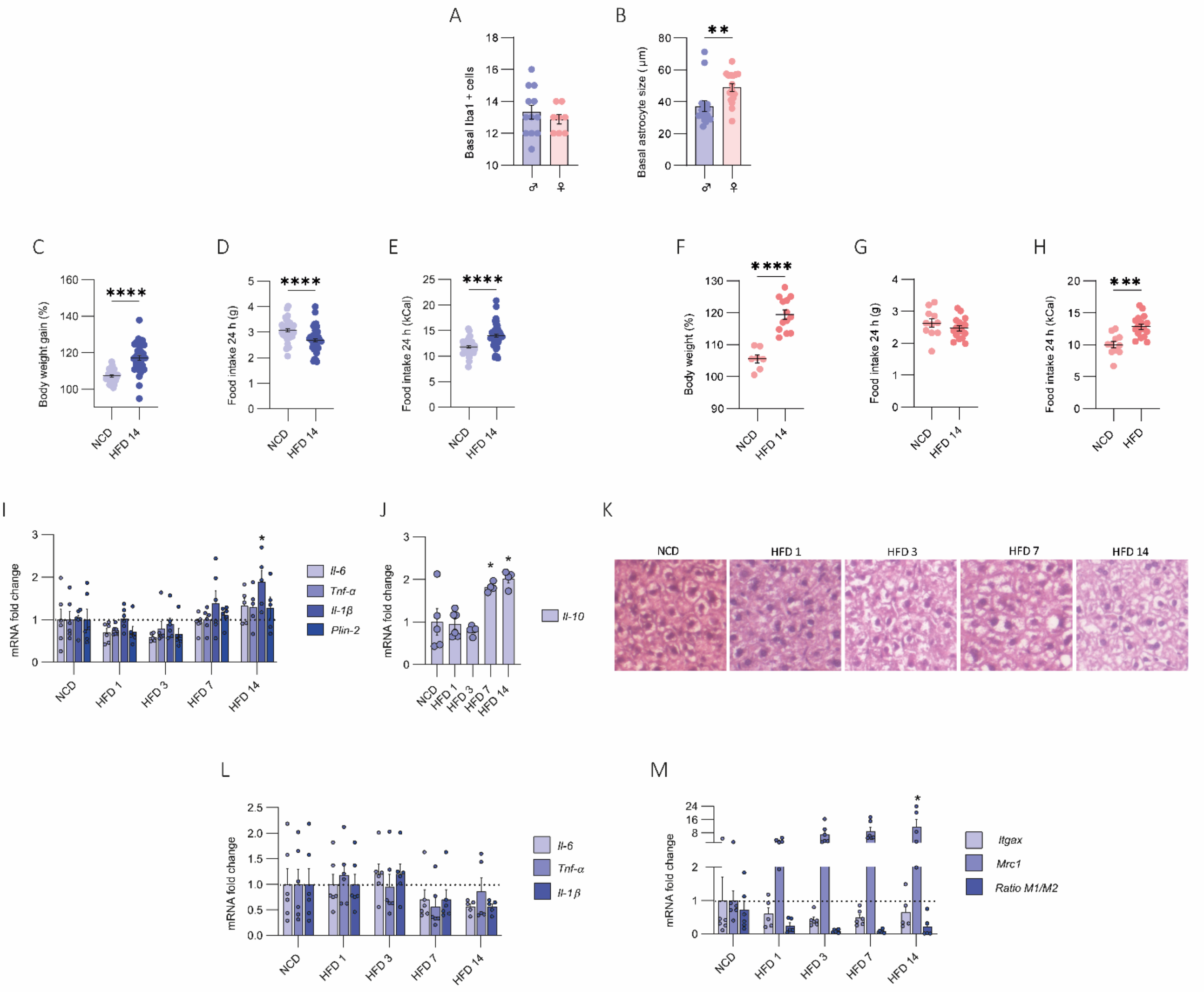


**Supplementary Figure 1 | Basal levels of microglia and astrocytes on the hypothalamus of male and female mice. Peripheral affections of the HFD.** (A) Basal number of Iba1+ cells comparing male and female mice. (B) Basal astrocytic size comparing male and female mice. (C-E) Body weight and food intake of male mice after 14 days of HFD feeding. n=21-23/group. (F-H) Body weight and food intake of female mice after 14 days of HFD feeding. n=7-13/group. (I) Analysis of the liver pro-inflammatory cytokines *Il-6, Tnf-α, Il-1β* and *Plin-2* mRNA levels by qRT-PCR of male mice. n=5-6/group. (J) Analysis of the liver anti-inflammatory cytokine *Il-10* mRNA levels by qRT-PCR of male mice. n=5-6/group. (K) Representative images of hematoxylin eosin staining in liver section of male mice fed with HFD. Images taken with a 10x objective. n= 5-6/group. (L) Analysis of the WAT pro-inflammatory cytokines *Il-6, Tnf-α and Il-1β* mRNA levels by qRT-PCR of male mice. n=5-6/ group. (M) Analysis of macrophages polarization analysis by *Itgax* for M1 and *Mrc1* for M2 mRNA levels by qRT-PCR of male mice. n=5-6/group. Data were represented as mean ± SEM. Statistical significance was determined by t-student or ordinary one-way ANOVA followed by Tukey's multiple comparisons post-hoc test. *p<0.05, **p<0.0.01, ***p<0.001, ****p<0.0001.

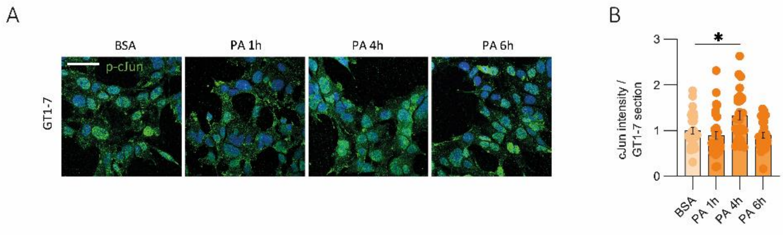


**Supplementary Figure 2 | c-Jun phosphorylation in GT1-7 after PA exposure** (A) Representative confocal images of p-cJun marker (green) in GT1-7 cells. (B) Quantification of p-cJun intensity in GT1-7 cells. Data were represented as mean ± SEM. Statistical significance was determined by ordinary one-way ANOVA followed by Tukey's multiple comparisons post-hoc test. *$p<0.05$. Scale bar 50 µm.

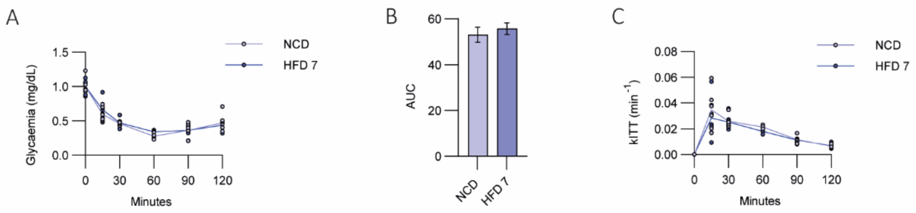


**Supplementary Figure 3 | Glycemia and kITT in male mice in response to intraperitoneal insulin administration after 7 days of HFD feeding.** (A) Glycemia in mg/dL. (B) Area under the curve of glycemia results comparing NCD versus HFD 7 days. (C) kITT calculation. n=7/group. Data were represented as mean ± SEM. Statistical significance was determined by t-student or two-way repeated measurement ANOVA followed by Sídák´s multiple comparisons post-hoc test.

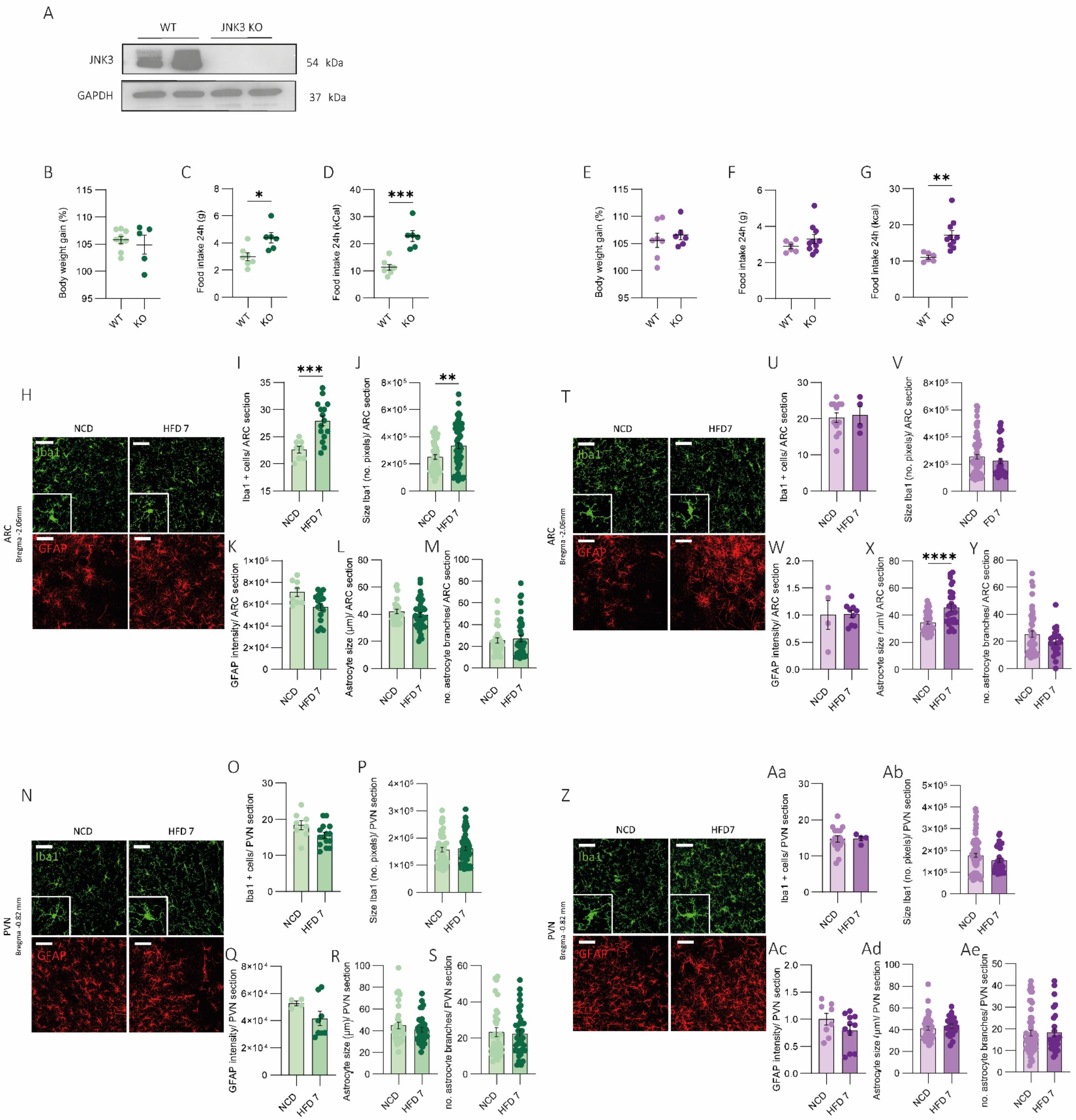


**Supplementary Figure 4 | Characterization of JNK3-KO mice after 7 days of HFD feeding.** (A) Representative Western Blot for JNK3 expression in hypothalamus from WT and JNK3 KO mice. (B-D) Body weight and food intake of male mice comparing WT versus JNK3 KO. n=5-7/group. (E-G) Body weight and food intake of female mice comparing WT versus JNK3 KO. n=5-7/group. (H,N,T,Z) Representative confocal images of Iba1 marker (green) and GFAP marker (red) in ARC and PVN sections of male and female JNK3 KO mice after 7 days of HFD exposure. Quantification of Iba1+ cells number of (I) ARC of male, (O) PVN of male, (U) ARC of female and (Aa) PVN of female. Quantification of microglia cell size by analyzing no. of pixels of (J) ARC of male, (P) PVN of male, (V) ARC of female and (Ab) PVN of female. Quantification of GFAP intensity of (K) ARC of male, (Q) PVN of male, (W) ARC of female and (Ac) PVN of female. Quantification of astrocyte cell size in µm of (L) ARC of male, (R) PVN of male, (X) ARC of female and (Ad) PVN of female. Quantification of no. of branches per cell of (M) ARC of male, (S) PVN of male, (Y) ARC of female and (Ae) PVN of female. n=3/group. Data were represented as mean ± SEM. Statistical significance was determined by t-student test. $*p<0.05$, $**p<0.0.01$, $***p<0.001$, $****p<0.0001$. Scale bar 50 µm.
